# BLAST : a short computerized test to measure the ability to stay on task. Normative behavioral data and detailed cortical dynamics

**DOI:** 10.1101/498691

**Authors:** Mathilde Petton, Marcela Perrone-Bertolotti, Diego Mac-Auliffe, Olivier Bertrand, Pierre-Emmanuel Aguera, Florian Sipp, Manik Batthacharjee, Jean Isnard, Lorella Minotti, Sylvain Rheims, Philippe Kahane, Vania Herbillon, Jean-Philippe Lachaux

## Abstract

This article provides an exhaustive description of a new short computerized test to assess on a second-to-second basis the ability of individuals to « stay on task », that is, to apply selectively and repeatedly task-relevant cognitive processes. The task (Bron/Lyon Attention Stability Test, or BLAST) lasts around one minute, and measures repeatedly the time to find a target letter in a two-by-two letter array, with an update of all letters every new trial across thirty trials. Several innovative psychometric measures of attention stability are proposed based on the instantaneous fluctuations of reaction times throughout the task, and normative data stratified over a wide range of age are provided by a large (>6000) dataset of participants aged 8 to 70. We also detail the large-scale brain dynamics supporting the task from an in-depth study of 32 participants with direct electrophysiological cortical recordings (intracranial EEG) to prove that BLAST involves critically large-scale executive attention networks, with a marked activation of the dorsal attention network and a deactivation of the default-mode network. Accordingly, we show that BLAST performance correlates with scores established by ADHD-questionnaires.

## 2. Introduction

The ability to maintain focused attention on a task until its full completion is crucial in many aspects of life, and has become a major topic of interest in several fields including education and professional sport (Kolb *et al*., 2011; Mangine *et al*., 2014; Caron, 2015; Diamond and Ling, 2016; Romeas *et al*., 2016). In most practical situations however, performance is impaired by Momentary Lapses of Attention or MLA (Weissman *et al*., 2006), which can last a few seconds or less (Peiris *et al*., 2006) during which cognitive resources are side-tracked towards processes unrelated to the instructed task (Smallwood and Schooler, 2006). MLA are pervasive in our daily life (Robertson *et al*., 1997; Killingsworth and Gilbert, 2010) and can have mildly negative to catastrophic effects from missing a crucial word in a complex explanation to hitting a bike rider while driving.

The behavioral characteristics and neuronal substrates of MLA remain to be established. In contrast, a related but somewhat distinct phenomenon - Mind-Wandering (MW) - has been the topic of extensive research lately, with clear evidence that the Default-Mode Network (Raichle *et al*., 2001) is critically involved (Mason et al., 2007; Christoff *et al*., 2009; Stawarczyk *et al*., 2011; Esterman *et al*., 2013; Poerio *et al*., 2017). But while MW usually causes the participant to lose complete sight of the general context of the task (s/he transiently « forgets » s/he’s supposed to perform a task), MLA are too short for task-sets (Sakai, 2008) to vanish completely, and mostly reveal themselves through isolated errors and/or a transient slowing down of reaction time (Weissman *et al*., 2006). Tasks used to study MW are not well-suited to study MLA : in popular designs such as the continuous performance test (CPT) or the Sustained Attention to Response Task (SART) (Bender *et al*., 2015; Phillips *et al*., 2015), participants are driven to automatize a specific behavioral response to frequent stimuli, while infrequent stimuli require a different response and failure to withhold the automated behavior in response to rare stimuli are indicative of MW. Such designs are well-adapted to study long and slow drifts away from the task - and their fMRI signature - but short MLA, which mostly occur during frequent stimuli and have little or no impact on automated behavior remain unnoticed.

We propose a new task specifically designed to detect MLA, and assess the ability to resist them and « stay on task », on a second-to-second basis : BLAST (Bron Lyon Attention Stability Task) measures the ability to activate selectively the right set of cognitive processes to continuously perform a short task with optimal performance, during one or two minutes, that is the duration of many task-units which require continuous and undivided attention (listening to a complex explanation, reading one or two pages of a book or playing a point in tennis). Such task-units are ubiquitous in daily life, and most longer tasks can be broken down into consecutive units of such duration, with little breaks in between. BLAST reveals fluctuations of attention within such units, with a temporal resolution on the order of one second; in that sense, it markedly differs from classic neuropsychology pen-and-pencil papers assessing attention only globally over the whole test. This is especially important as the variability of reaction times is increasingly used as a measure of attention instability for the diagnosis of attention deficit hyperactivity disorder (ADHD) (Tamm *et al*., 2012; Gmehlin *et al*., 2016).

The paper combines three studies in one : a) an in-depth analysis of behavioral data in a large population of children and teenagers (>800) which reveals the main factors impacting attention stability (age, sex, ADHD-scores), b) normative behavioral data stratified by age from a larger (>6000) population aged 8 to 70 and c) a detailed analysis of the large-scale and highly dynamic cortical network supporting BLAST, from the most precise neural recordings available in humans (intracranial EEG).

## 3. Participants and materials

### 3.1. Behavioral study

Behavioral data were collected in schools (SCHOOL database) from 859 participants (446 females, 103 left-handers), aged 8 to 18, all free of psychiatric/neurological disorders and with an appropriate behavior during the test (silent, showing no obvious lack of motivation). Participants performed the task on iPads wearing headphones and facing a wall with no view of each other, in groups of one to four, in a large classroom. Sessions lasted one hour to include BLAST, plus several variants of the main task (see tasks description). The study was approved by a national ethical committee (IRB00003888 - FWA00005831).

In addition, a larger normative sample was collected within a science exhibit (« expo cerveau » - cité des sciences de la Villette, Paris) where BLAST was proposed as an interactive test to evaluate the visitors’ ability to resist endogenous distractions. The test comprised a series of 30 trials, after a learning stage of 15 trials. Immediately after task completion, visitors indicated their age, gender and whether they had been distracted by external events. The MUSEUM database consists of 6954 participants under 70 (4049 females) reporting no external distractions (74% of 9446 visitors).

### 3.2. Electrophysiological study

Intracranial EEG recordings (iEEG) were obtained from 32 patients (19 women, mean age: 28.7, Std: 8.6) candidate for epilepsy surgery. Two patients were not able to perform the task and were excluded from the analysis. Participants were stereotactically implanted with multilead EEG depth electrodes following the classic clinical procedures of the neurological hospitals in Grenoble and Lyon. They had previously written informed consent to participate in the study and experiment have been approved by the local ethical Committee of Grenoble Sud-Est V (Study 0907 - ISD et SEEG, CPP 09-CHUG-12).

Five to seventeen semi-rigid electrodes were stereotactically implanted in each participant. Each electrode was a linear array with a diameter of 0.8 mm, and included between 5 and 18 contact leads along its shaft (2mm wide, with a 3.5 mm center-to-center spacing between consecutive leads) (DIXI Medical, Besançon, France). The precise anatomical location of each lead was measured from patient’s individual pre-implant MRI, co-registred with post-implant MRI using an in-house BrainVisa plugin (IntrAnat), that enables the trajectory reconstruction of precise models of the SEEG electrodes onto post-implant MRI (see BrainVisa, brainvisa.info).

Intracranial EEG (iEEG) was recorded with standard 128-channels and 256-channels iEEG acquisition system (Micromed, Treviso, Italy), with a reference in the white matter. Data were bandpass-filtered online [0.1-200 Hz] and sampled at a minimal frequency of 512 Hz, then re-referenced offline by subtracting for each site the signal recorded 3.5 mm away on the same linear electrode (the nearest neighbor : bipolar montage).

Signals were visually inspected with an in-house, freely downloadable software package for electrophysiological analysis (ELAN-pack) (Aguera *et al*., 2011) to reject leads with epileptiform activity. High-Frequency Activity [50-150 Hz], or “high-gamma activity”, was extracted with in-house scripts in Matlab (the Mathworks, inc) following our usual procedure (Ossandón *et al*., 2012). That procedure can be summarized as follows: bipolar iEEG signals are first bandpass-filtered in ten consecutive 10-Hz-wide frequency bands (50-60 Hz, 60-70 Hz,…, 140-150 Hz). Each of those ten bandpass-filtered signal is then transformed into its envelope using a standard Hilbert transform and each envelope is then expressed in % of its mean value (as calculated across the entire experiment). In other words, each envelope time series is simply divided by its mean across the entire recording session and multiplied by 100. In the last step, the ten normalized envelope signals (in %) are averaged together (across the ten frequency bands) to provide one single time-series: the High-Frequency Activity [50 to 150 Hz], or HFA. Significant stimulus-induced HFA variations are detected statistically by comparing HFA for each time sample [between 0 and 3000 ms relative to the first stimulus of a trial] with the mean HFA during a [-200 ms : 0 ms] prestimulus baseline (Wilcoxon signed rank test corrected with a threshold p = 0.01). HFA measured in specific windows following the initial stimulus (e.g. [100 ms : 200 ms post stimulus onset]) is compared across experimental conditions using nonparametric Kruskall-Wallis tests (p<0.01).

Dynamic, group-level, visualizations of task-induced HFA variations onto the standard Montreal Neurological Institute (MNI) single-subject brain were performed with HiBoP (Human Intracranial Brain Observations Player), a new iEEG visualization software developed in our laboratory within the Human Brain Project, which will be released freely in the coming year.

### 3.3. tasks description

#### 3.3.1. BLAST

BLAST repeatedly asks participants to find a target letter (the Target) in a subsequent two-by-two array of four letters (the Array), with new letters every trial (Target and Array) (Fig.1). Each trial starts with the presentation of the Target for 200 ms, followed by a mask (#) replaced after 500 ms by the Array (which stays on screen until the manual response). The next trial starts after a pause of 800 ms if the previous response was correct (with a central fixation symbol #), 4800 ms if it was incorrect (which corresponds to a time penalty of 4000 ms) and 3800 ms if the participant did not respond. The penalty system encourages participants to respond only if they are sure of their choice. Experiments were run on a tactile tablet for the SCHOOL and MUSEUM populations: participants were instructed to hold both hands slightly above the screen, on each side of the letter display, and give a gentle touch with their dominant hand (resp. non-dominant hand) when the target was absent (resp. present). During iEEG recordings, patients performed BLAST on a PC synchronized with the EEG acquisition system (The experiment was performed using Presentation^®^ software (Version 18.0, Neurobehavioral Systems, Inc., Berkeley, CA); they pressed gamepad buttons with their left or right index finger (using the same “dominant/non-dominant hand”-“no/yes” mapping as participants of the SCHOOL and MUSEUM populations). Letters were presented foveally in black on a light gray background. Motor responses were immediately followed by a 50 ms auditory beep signal indicating success or failure (one sound for each outcome), at comfortable hearing level. The sequence of « target present » and « target absent » trials was pseudorandom, with the same number of trials for both types. Performance was measured on a total of 30 trials, for a total duration around one-minute (depending on reaction times).

**Figure 1:**
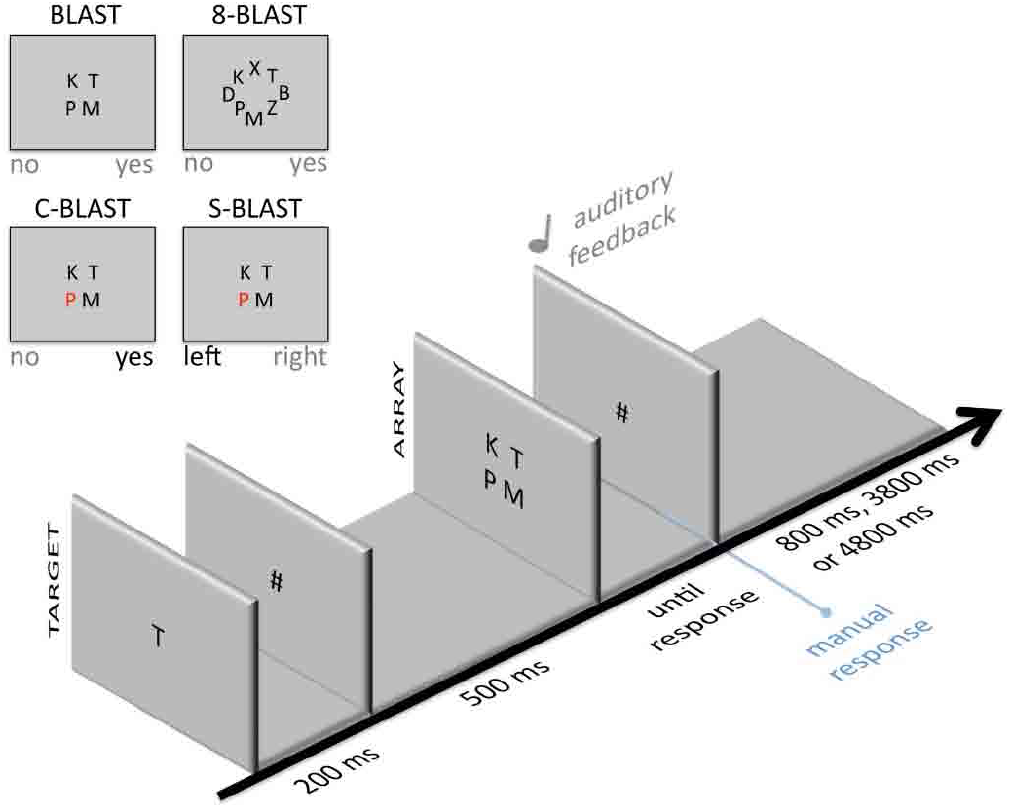
General design of BLAST and its variants. The lower panel shows the general structure of a trial with the sequential display of the Target and the Array, and new letters every trial. The duration of the inter-trial mask depends on the participant’s response (800 ms when the response was correct, 4800 ms if it was incorrect, and 3800 ms when the participant did not respond). The upper panel shows the specificities of the three main variants of BLAST in the case of a left-handed participant: the only difference with BLAST is the Array screen, with eight letters in 8-BLAST instead of four, four letters in C-BLAST with a salient color for the target when it is present, and four letters in S-BLAST with a red letter every time, indicating which side of the screen must be touched (the same as the red letter).

The global instruction was to settle in a steady and reasonably fast pace while avoiding errors (with an explicit analogy to car drivers who avoid accidents at all costs, but nevertheless move forward at a decent speed). The time penalty was precisely to discourage risky strategies, and to minimize the distraction caused by errors and negative feedbacks on subsequent trials: participants were given time to refocus.

#### 3.3.2. Variations of BLAST: adding and removing cognitive components

Participants of the MUSEUM database performed BLAST after a separate training phase of 10 trials. Participants of the SCHOOL database were tested during a one-hour session which included BLAST, plus several additional tasks (C-BLAST, S-BLAST and 8-BLAST, Table.1) defined as follows (Fig.1 and Table 1): C-BLAST (“Color-BLAST”) was a simplified version of BLAST, with no memory or visual search component: the Target letter was displayed in red in the Array to pop out when it was present (other letters were black). The instruction was simply to indicate manually if one of the letters in the Array was red (« no » with the dominant hand, and « yes » with the other hand, as in BLAST). S-BLAST (“Side-BLAST”) pushed the simplification one step further, with a direct stimulus-response contingency: a singleton red Target was present in all Arrays and participants simply had to touch the screen on the same side as the Target (simple reaction task). In contrast, 8-BLAST was a more difficult version of BLAST with a circular eight-letters Array instead of the square four-letters Array (longer visual search). Trials in all tasks had similar timing parameters (as described earlier).

C-BLAST, S-BLAST and 8-BLAST were used to evaluate the cost and benefit (in reaction time) of adding or removing critical cognitive components to BLAST. They were performed during a specific session following an adaptive design (see next paragraph). The description of that design, and the presentation of the corresponding results can be found in the next section (“Adaptive version of BLAST”).

**Table 1.** versions of BLAST used for each subpopulation

| Population | Versions of BLAST |
| --- | --- |
| SCHOOL | BLAST, C-BLAST, S-BLAST, 8-BLAST |
| MUSEUM | BLAST |
| SEEG | BLAST, C-BLAST, S-BLAST |

#### 3.3.3. Adaptive versions of BLAST

In schools and during iEEG sessions, C-BLAST, S-BLAST, 8-BLAST (and BLAST, in addition to the standard design described above) were performed in an adaptive, staircase, design, in which participants were asked to achieve a series of five consecutive successful trials, under increasing time pressure. Trials of each task were designed as in BLAST, with the presentation of the Target (200 ms), followed by the mask (700 ms) and the Array until the participant’s motor response (i.e. the same timing as BLAST). The overall task design differed from BLAST, however, in order to evaluate how fast a participant could respond under such an accuracy constraint. In the adaptive design, the explicit objective was to achieve series of five consecutive correct trials, faster than an adaptive time limit which decreased with each successful series, until the participant could no longer succeed. We measured TIME_LIMIT, the best limit s/he could reach and the median of all correct reaction times MEDIAN_RT_CORRECT. The time limit was initialized to 10 seconds, then updated after each successful series to the slowest reaction time of that series, minus a 20 ms decrement (to converge as fast as possible to the participant’s limit): the participant was challenged to achieve a slightly faster series. Series were immediately interrupted after any response incorrect or too slow (relative to the ongoing time limit), and the new time limit was simply increased by a 20 ms step for a new, easier, challenge. The objective (five trials) is obviously arbitrary: it is a compromise between a goal that is too hard (and would discourage participants) and one that is too easy (and can easily be achieved by chance).

### 3.4. Post-test questionnaires

Participants of the SCHOOL population filled up a questionnaire immediately after the experiment to rate several subjective factors including i) their motivation during the task (MOTIV, on a 1 to 10 scale from ‘not motivated at all’ to ‘extremely motivated’); ii) if they had to focus during the task (FOCUS, on a 1 to 10 scale); iii) how stressed they felt (STRESS, on a 1 to 10 scale); iv) if they felt they had been mind-wandering during the task (MW on a 1 to 4 scale, from “not at all” to “a lot”), and v) if they had been distracted by external noise (NOISE, on a 1 to 4 scale, from « never » to « very often”).

Another questionnaire, filled at home by parents, asked about various aspects of the participants’ life outside of school, focusing on TV and video-game usage (number of hours) and hobbies (sport and artistic activities). Finally, parents also filled an ADHD rating scale IV home version (DuPaul *et al*., 1998) based on the global behavior of their son or daughter the past 6 months. This questionnaire includes 18 questions addressing separately two dimensions (inattention and hyperactivity/impulsivity); it assesses symptoms of attention deficit/hyperactivity disorder (ADHD) according to the diagnostic criteria of the Diagnostic and Statistical Manual of Mental Disorders (DSM IV).

### 3.5. Behavioral indices

During BLAST, reaction time (between the onset of the Array and the response) and accuracy were measured for 30 consecutive trials. Attention stability indices were computed from those measures. Our primary intention was to reveal the propensity to undergo momentary lapses of attention (MLA, or conversely, momentary “peaks of attention”). It led us to depart from standard measures such as the variance of the entire reaction times series, which fails to capture the moment-to-moment dynamics of attention (since the variance might be the same for a participant alternating between highly focused and highly distracted episodes, and another participant mildly focused throughout the task). Clearly, new indices had to be designed to integrate the number, the duration and the local stability of attentive episodes. It is not uncommon to propose several scores for the same test to measure slightly different cognitive components (Christidi *et al*., 2015). In the following, we describe and motivate the indices we derived from BLAST for neuropsychological assessment. All indices can be adapted to any task with constant difficulty which measures reaction times repeatedly and steadily.

#### 3.5.1. INTENSITY

The first behavioral measure, INTENSITY (figure 2), derives from the natural assumption that highly focused individuals tend to respond fast and with few errors. In other words, a performance graph showing reaction times (y-axis) for all the trials (x-axis) should display long series of hits below a given time limit T (horizontal line), even when T is low (see Figure 3, bottom panel on the left). If we cumulate the length of all successful series below T (i.e. with a reaction time consistently faster than T), we reach a measure P(T) which should be high for “highly-focused” individuals even when T is small (i.e., 500 ms). To penalize errors and to be consistent with the adaptive designs which required participants to generate successful series of five trials (in reference to the description of “adaptive version of BLAST” above), we computed a measure P(T), which included only series longer than five consecutive wins, cumulating N-5 « points » for every such series (where N is the length of that series). P(T) is the number of points for time limit T; it is computed for every value of T between 300 ms and 1500 ms (no one could generate series of hits faster than 300 ms). That scoring system penalized errors, as any error interrupted the ongoing series and the participant had to succeed at least five new times before her score increased again. Graph P(T), which increases with longer T, immediately distinguishes visually between several types of participants/strategies (Fig.3) : fast reaction times with many errors (‘globally impulsive participants’), slow reaction times with few errors (‘globally meticulous’), slow reaction times with many errors (‘slow inattentive’ type) and fast reaction times with few errors (‘fast and focused’). Fast and focused participants are characterized by a large area under the curve P(T): for that reason, we defined a behavioral indice of attention, INTENSITY = 100xAUC(P)/max(AUC), where AUC is the area under the curve P(T) and max(AUC) its maximal theoretical value. INTENSITY refers to, and is high for, the “fast and focused» type. One might object that lower values of INTENSITY might not disambiguate the ‘impulsive’ and the ‘meticulous’ types, but those two profiles can be differentiated by considering also the median of their reaction times (and graph P(T)).

**Figure 2:**
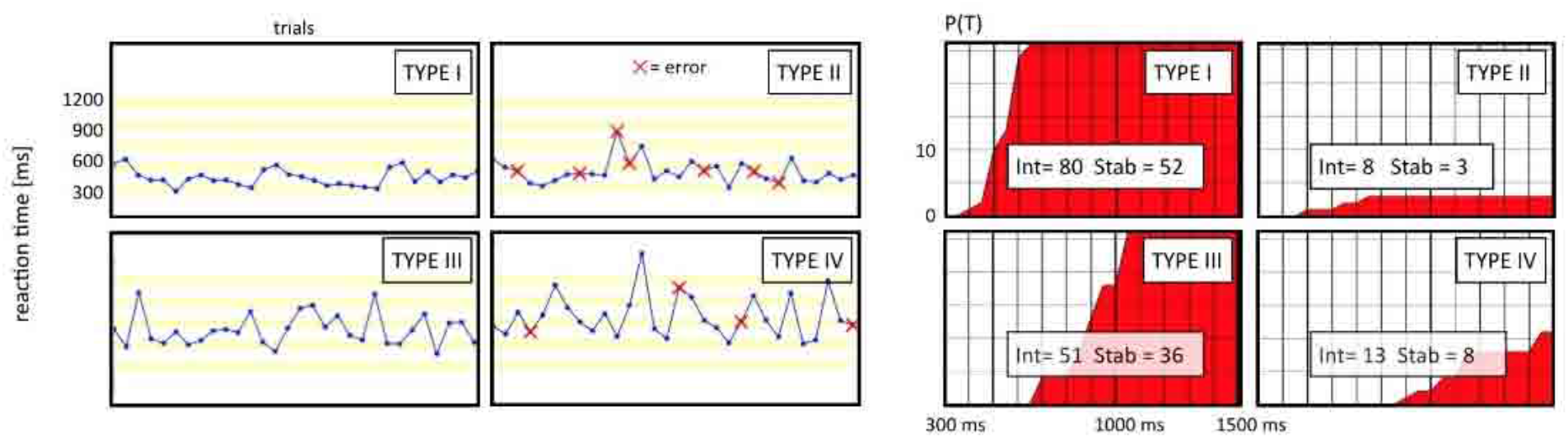
Scoring systems for INTENSITY and STABILITY. INTENSITY and STABILITY are computed from graphs P(T) and S(T’) respectively (bottom and top right panels, respectively), from the ratio of the area under the curve (red) divided by the total area of the square englobing that curve (white). The bottom left graph illustrates the calculation of P(T) from the reaction times for an example value of T (600 ms : 9 points) (« instantaneous » standard deviations of reaction times - computed over three consecutive trials, are shown as gray vertical bars). The procedure is repeated for all T between 300 ms and 1500 ms to generate red plot P(T). The top left graph illustrates the calculation of S(T’) from the « instantaneous » standard deviation s(trial) of the normalized reaction times (r’(trial), see methods) for an example value of T’(20 : 12 points). The normalized reaction times r’(trial) are simply reaction times expressed as % of the median reaction time. S is depicted in the top left graph as vertical bars (green = s less than 10; yellow = s less than 20; red otherwise, note that s is set to 40 for every error) and red vertical bars in the upper left graph). The procedure is repeated for all T’ between 0 and 40 to generate red plot S(T’) on the right.

**Figure 3:**
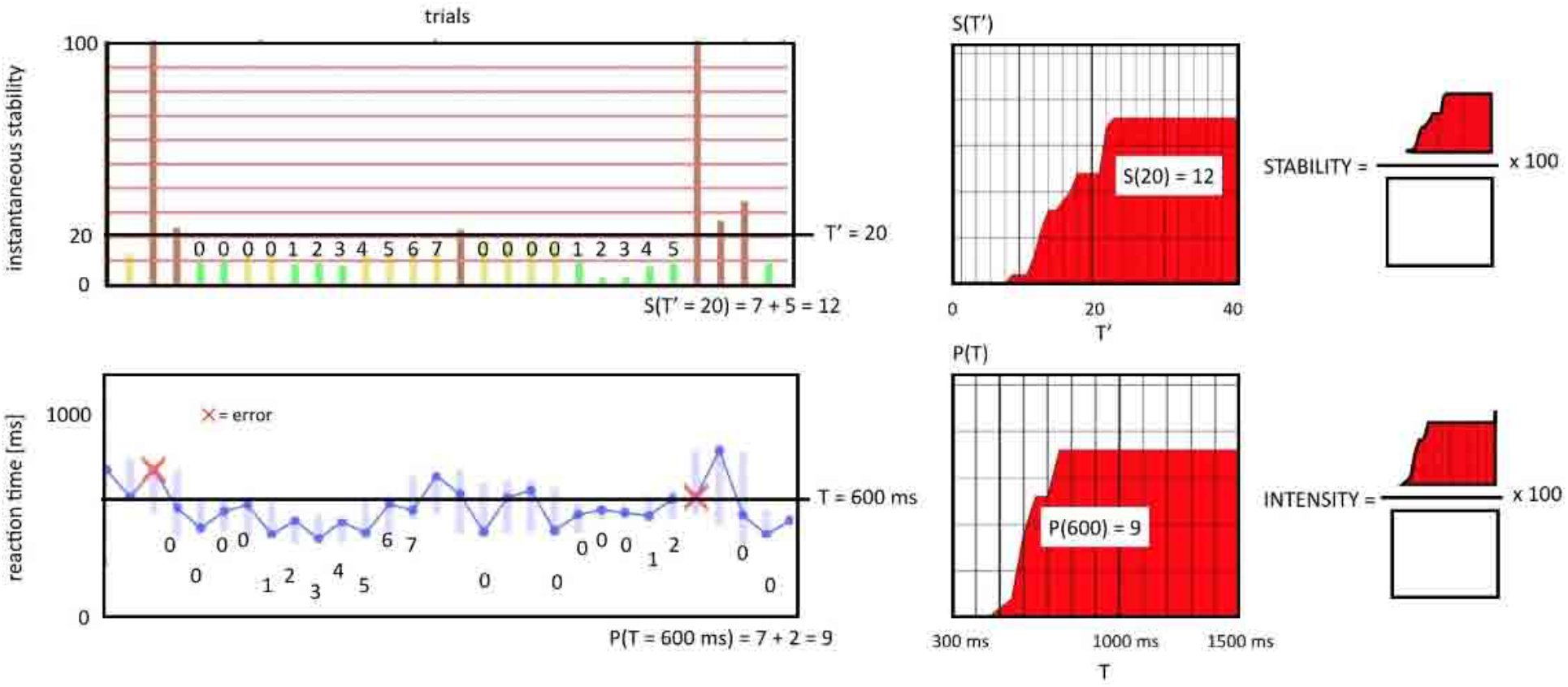
Examples of response profiles for BLAST. The left panel displays reaction times and errors for four participants (same age-group : 16-17 year-old), which correspond to four different response-types (right panel). Type I is fast, steady and accurate and exemplifies what we called FAST AND FOCUSED individuals. P(T) (right panel) is high for short reaction times and quickly reaches maximal values: most of the square is red. Type II is as fast as Type I, but has many errors (IMPULSIVE type) : consequently, P(T) mostly colors the bottom end of the square, from fast to slow reaction times. Type III corresponds to the METICULOUS type, with very few errors but slow reaction times. P(T) mostly colors the right end of the square. Finally, Type IV corresponds to the SLOW INATTENTIVE type, with many errors and slow reaction times. Neuropsychologists might derive a parallel typology from STABILITY graphs.

##### 3.5.2. STABILITY

One limitation of INTENSITY is that it is mathematically lower for participants with slower reaction times (typically, elderly people). Yet, it is clear that someone with slow but close to constant reaction times, and no errors, is fully « on-task ». Therefore, INTENSITY should be complemented with another measure which emphasizes stability more than speed: this is STABILITY(figure 2). STABILITY was derived from the assumption that task-irrelevant cognitive processes add “noise” to reaction times (Gmehlin *et al*., 2016): it was thus computed from the “instantaneous” stability of reaction times r(trial i). And since our intention was to minimize the effect of the overall speed, we considered the stability of “normalized reaction times”; i.e. reaction times expressed in % of the median reaction time for the entire task: r’(trial i) = 100*r(trial i)/median(r(all trials)). We defined the instantaneous stability of attention for trial i as instab(trial i) = std(r’(trial i-1):r’(trial i+1)), it is computed over sliding windows of three consecutive trials for maximal temporal precision. STABILITY was computed from instab(trial i) following a procedure similar to the computation of INTENSITY from r(trial i). To penalize errors as in INTENSITY, instab was set to a maximal value (40) for unsuccessful trials. As with INTENSITY, we devised a scoring system S(T’) which accumulates N-5 points for each series of N>4 winning trials for which instab stays below T’ (a procedure repeated for T’ values between 0 and a maximum of 40). Graphically, instab measures the width of the « tube » in which the reaction time plot seems to be confined locally (i.e. for three consecutive trials) (Fig.2), and S(T’) integrates the length of such series, excluding series shorter than 5 for the same reason as in INTENSITY. In short, STABILITY evaluates the ability to generate long and narrow « tubes » of reaction times: STABILITY = 100xAUC(S)/max(AUC), where AUC is the area under the curve S(T’) and max (AUC) its maximal theoretical value.

##### 3.5.3. Other behavioral indices

In addition to INTENSITY and STABILITY, we considered more standard indices such as the overall duration of the task (including time penalties and divided by the number of trials: MEAN_DURATION), the percentage of incorrect trials (PCT_ERRORS), the median and the standard deviation of the entire series of reaction time (excluding time penalties: MEDIAN_RT and STD_RT). Finally, upon the repeated request of teachers helping with data collection, we designed a measure of the duration participants actually spend “on-task” during BLAST (expressed in % of the task duration). Assuming again that episodes spent “on-task” are characterized by stable and accurate responses, we proposed a measure based on instab(trial i): FOCUS. FOCUS is the percentage of trials for which instab(trial i) is less than 20 (remember that instab is set to 40 for unsuccessful trials). We are fully aware that the cut-off value of 20 is arbitrary: it was chosen after numerous attempts to relate instab with self-reports of how participants felt. A special version of BLAST was designed to interrupt the task at any moment and ask participants about their attention. They felt reasonably on task with the feeling of being focused when instab was less than 20. Although debatable, a choice had to be made and alternative cut-off values (i.e., 10) resulted in measures highly correlated with FOCUS.

#### 3.6. Statistical analysis

Unless specified otherwise, the effect of a given factor (e.g. AGE) was tested using a non-parametric Kruskal-Wallis test (abbreviated KW), as Kolmogorov-Smirnov (KS) test revealed general nonnormality of our indices. For some analysis, carefully identified in the result section, indices were first normalized within each age group (Z-score normalization: substrate the mean value for that particular age group and divide by the standard deviation), to study the effect of a factor independently of age (e.g. GENDER). This normalization is explicitly mentioned in the text each time it is used.

### 4. Results

#### 4.1. Behavioral results

We tested several major predictions: i) that the ability to stay-on-task would increase from children to teenagers and adults, with a plateau in the early twenties at the end of the maturation of the prefrontal cortex and the executive system (Gogtay *et al*., 2004; Toga *et al*., 2006); ii) that children and teenagers with high inattention scores (according to the ADHD questionnaire) would perform worse than participants matched in age with lower inattention scores, and iii) that attention might be affected negatively or positively by extra-curricular activities (TV and video-games, art, sport) and by sociodemographic factors (e.g., education level of the parents). The SCHOOL database was designed to test those predictions, as it combined BLAST behavioral measures with questionnaires filled by parents. In addition, short questionnaires filled by participants at the end of the test allowed us to evaluate whether they had performed the test in good conditions (motivated, with no external distractions) and considered actually BLAST as attention-demanding.

##### 4.1.1. Analysis of post-test questionnaires (self-reports)

BLAST was designed to be attention-demanding and yet pleasant, although not to the degree reached by video-games carefully designed to capture and hold the player’s attention. Participants confirmed we reached our goal: a large majority enjoyed BLAST and said they had to focus (MOTIV, “did you enjoy the test? »: 92% above 5, 64% above 7.5 on a 0 to 10 scale; FOCUS, “how focused were you? »: 92% above 5, 69% above 7.5 on a 0 to 10 scale). They also reported little or no distraction by external noises (NOISE: “were you disturbed by external noise? »: 95% below 2 on a 1 to 4 scale, i.e. « never » or « rarely »), which means that our experiment was well-suited to study *endogenous* distraction, as intended.

##### 4.1.2. BLAST scores and ADHD rating scale

A standard ADHD-questionnaire was collected from 692 participants (SCHOOL database), with nine questions related to inattention specifically and nine questions related to hyperactivity (DuPaul *et al*., 1998). Parents were asked to rate each statement between 0 and 3 (from 0 - « almost never » - to 3 - « very often”), leading to a global inattention score (for nine questions, from 0 to 27), a global hyperactivity score (0 to 27) and a global ADHD score (the sum of the two). We tested for relationships between each global score and each individual question, and all behavioral indices.

To test for possible relationships between global scores and BLAST indices (e.g., INTENSITY,…), BLAST indices of each participant were transformed into a group letter (A, B or C), indicating whether that participant was in the upper, middle or lower 33% of her age group for that particular index (performance-wise). We tested the Ho-hypothesis that global scores did not differ across groups (A,B,C) and rejected that hypothesis for the global INATTENTION score (KW, p<0.005) for all indices (STABILITY, INTENSITY, FOCUS, MEAN_DURATION, MEDIAN_RT, PCT_ERRORS and STD_RT) : participants had a higher inattention score in groups (i.e., letters) with the poorest performance. In contrast, no significant difference between groups were found when considering the global Hyperactivity score. The global ADHD score, which adds the global inattention and hyperactivity scores, differed across groups when considering indices INTENSITY, FOCUS and MEAN_DURATION (KW, p<0.01).

A more detailed question-by-question analysis revealed a finer pattern. Since each question was associated with a rating between 0 and 3, we tested for an effect of that rating on normalized BLAST measures (Z-score, relative to age-group, as described in the methods section). We found a significant effect of the rating for 5 of the 9 questions related to Inattention : question 1 (effect on STABILITY, INTENSITY, FOCUS, MEAN_DURATION, MEDIAN_RT and STD_RT, p<0.005) [« Fails to give close attention to details or makes careless mistakes in schoolwork»]; question 3 (for all indices also, p < 0.001) [«Has difficulty sustaining attention in tasks or at play »],; question 11 (p<0.005 for all indices, except STD_RT) [«Avoids tasks (eg, schoolwork, homework) that require a sustained mental effort»]. Question 9 had only an impact on STABILITY (p<0.01) [«Has difficulty organizing tasks and activities»] and Question 7 on STABILITY and STD_RT [«Does not follow through on instructions and fails to finish work »]. We found no significant effect for the four remaining questions, which were less directly related to the specifics of BLAST (as BLAST involves very clear task instructions, a short duration and a quiet environment): « Is easily distracted by outside stimuli »; « is forgetful in daily activities »; « does not seem to listen when spoken to directly »; « Loses things necessary for his tasks or activities ».

The same analysis of hyperactivity questions led to significant results for one question only [« interrupts or is intrusive »]: STABILITY and PCTERRORS, KW, p < 0.01). Overall, our analysis indicates that BLAST is almost uniquely sensitive to the Inattention component of the ADHD rating scale, and more precisely with the observed inability to stay-on-task.

##### 4.1.3. BLAST scores and participants daily conditions and occupations

Using the same approach as above, we tested for a possible effect on BLAST of several important elements of children’s life, including the socio-professional category of their parents (mother/father, separately) and extra-curricular activities (time spent watching TV or video-gaming, practicing sport or artistic activities). We found no significant effect on any of the BLAST indices (normalized by age). We also tested whether performance was affected by the time of day at which BLAST was performed (early or late morning, early or late afternoon), but found no significant effect on any of the BLAST measures.

##### 4.1.4. Normative data and age and sex effects

Normative data were computed from the MUSEUM database and displayed graphically for all behavioral indices in Fig.4 in a form that reveals a clear effect of age. All performance indices were affected by age, including all measures of attention (Kruskal-Wallis, p< 10^−22^ for all), with a systematic increase of performance with age for participants younger than 25. Participants also reported their gender, and we found significant differences between the two genders on several behavioral indices (Kruskal-Wallis, p<0.005 for INTENSITY, PCT_ERRORS an MEAN_DURATION), even when considering normalized scores (relative to same-age participants, see methods). Females outperformed males with a more stable attention and less errors.

**Figure 4:**
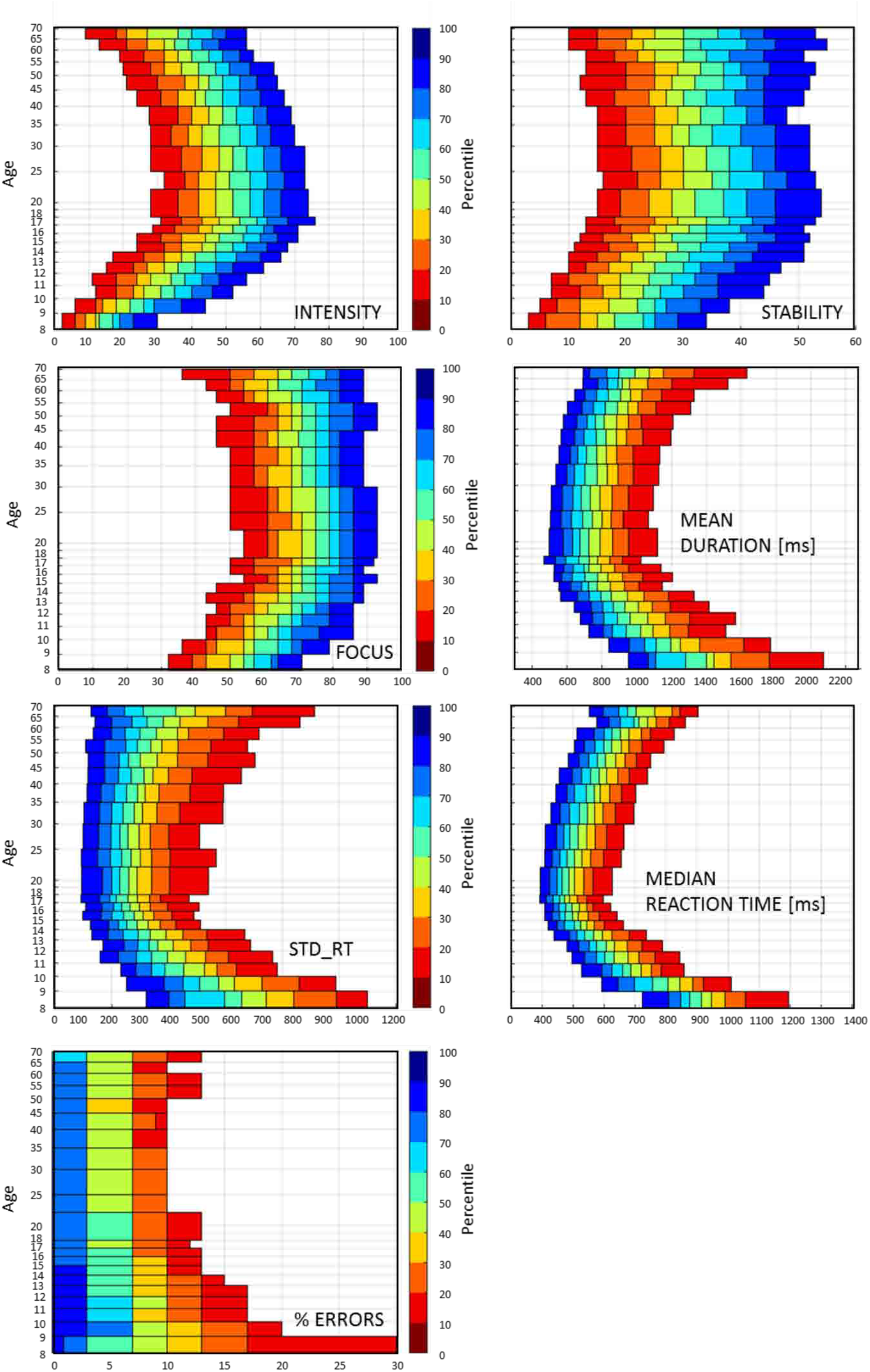
Age-stratified normative data for the main behavioral indices quantifying the ability to stay on task in BLAST. A neuropsychologist using the graphs to interpret BLAST scores, should identify the age of the participant on the y-axis and his/her score on the x-axis, then use the color-code to evaluate immediately the proportion of participants of the same age with a lower score (90,80,70 %…).

The pervasive influence of age led us to compute and display nine quantiles (10 to 90%, by step of 10%) as a function of age, with a one-year stratification between 8 and 18. A neuropsychologist using the graphs to interpret BLAST scores, should identify the age of the participant on the y-axis and her score on the x-axis, then use the color-code to evaluate immediately the proportion of participants of the same age with a higher score (90,80,70 %…). Supplementary Fig.1 also show polynomial functions modelling the age effect, where the dependence of quantile values on age was nicely fitted by third-order polynomial functions q(age) = a.y3 + by2 + cy + d, where y = log(age) (polynomial fitting in MATLAB interface, the Mathworks, inc.).

It is worth noticing that none of the distributions is bimodal, for any age group. The distributions of all behavioral indices have only one peak, and any attempt to define a cutoff score to identify impairment (“high” vs “low” attention scores) would be arbitrary and debatable.

##### 4.1.5 Test-Retest

Test-retest reliability was evaluated from data of 122 participants who performed BLAST twice, using two measures: the Pearson’s correlation coefficient (Rho) and the intraclass correlation coefficient - (icc; irr package of R with parameters: ‘oneway’ and ‘consistency’). Data are shown in Table 2; unsurprisingly, the most reliable measure is the median reaction time (Pearson Rho = 0.93, icc = 0.90), while most measures reach 0.6 or higher with both methods, with a maximum reliability for INTENSITY (Rho = 0.8, icc = 0.81) and a minimum reliability for MEAN_DURATION (Rho = 0.49, icc = 0.49). These values are in line with test-retest reliability measures of standard tests of executive functions (Lowe and Rabbitt, 1998).

**Table 2:** Test-retest reliability for the main behavioral indices of BLAST performance.

|  | **Icc (all)** | **R pearson (all)** |
| --- | --- | --- |
| INTENSITY | 0.80 | 0.81 |
| STABILITY | 0.65 | 0.65 |
| FOCUS | 0.52 | 0.55 |
| MEAN_DURATION | 0.49 | 0.49 |
| STD-RT | 0.60 | 0.61 |
| MEDIAN_RT | 0.90 | 0.93 |

##### 4.1.6. Comparison of BLAST and CODE

Thirty-seven participants of the same age group (16-17) performed both BLAST and a standard pencil-and-paper test of sustained attention (CODE), a subtest of processing speed index from the Wechsler Intelligence Scale for children and the Wechsler Adult Intelligence Scale respectively WISC-IV and WAIS-IV (Wechsler, 2003, 2008). CODE consists in matching symbols to numbers as quickly as possible during two minutes, according to written instructions. We found significant correlations between CODE (number of correct responses) and most behavioral indices of BLAST: INTENSITY (Spearman Rho Correlation Coefficient = 0.57, p = 0.0002); STABILITY (Rho = 0.48, p = 0.002); MEAN_DURATION (Rho = −0.59, p < 1e-5: participants better at CODE were faster at BLAST); FOCUS (Rho = 0.61, p < 1e-5); MEDIAN_RT (Rho = −0.55, p = 0.0004); PCT_ERRORS (Rho = − 0.39, p < 0.05 - less errors at BLAST for higher scores at CODE).

##### 4.1.7. Adaptive versions of BLAST

The adaptive versions of BLAST measured the participants’ fastest reaction time under an accuracy constraint (five consecutive successful trials, see methods). Unsurprisingly, we found that such reaction time limit was longer for more complex tasks. The median reaction time (for correct trials) increased from S-BLAST to C-BLAST, BLAST and 8-BLAST (see Supplementary Fig. 2): S-BLAST (MEDIAN_RT_CORRECT, mean = 300 ms +/− 60 ms); C-BLAST (mean = 390 ms +/− 90 ms);

BLAST (mean = 538 ms +/− 105 ms); 8-BLAST (mean = 687 ms +/− 148 ms). The cost of increasing complexity was higher for younger participants but on average, the cost of increasing search difficulty - from four to eight items - and the benefit of removing the search component - from four items to one item - were roughly equivalent to 150 ms (BLAST to 8-BLAST, + 149 ms; BLAST to C-BLAST, −148 ms). The cost of converting the abstract answer (yes or no) into a motor response (S-BLAST to C-BLAST) was 90 ms.

##### 4.1.8. One minute or ten minutes?

Considering that BLAST lasts only about one minute - in its thirty-trials version - one might question its validity to assess the ability to stay on task for longer durations, over segments of ten minutes for instance. One might argue that individuals who can remain sharply focused for one minute are not necessarily the same who can perform steadily for ten minutes. We addressed this question with a database of 121 participants aged 10-12 (65 males) performing a 200 trials version of BLAST (separate from the SCHOOL and MUSEUM datasets and participants). We computed our main indices over the first 30 trials and correlated their values with the same indices measured over the entire session (200 trials), Table 3 contains the result, as well as the same correlation analysis when considering the first 40, 50 and 60 trials. We found that for several indices (INTENSITY, MEAN_DURATION and MEDIAN_RT), the correlation (Pearson’s correlation coefficient) was already high for 30 trials (0.75, 0,75 and 0.87 respectively), meaning that the ability of an individual to be fast and focused during ten minutes could be fairly well estimated in one minute only. The full analysis suggests that the STABILITY and FOCUS indices over ten minutes should rather be estimated with a 50 trials version of BLAST (R = 0.78 and 0.67 respectively). The conclusion that long-term (10 minutes) attention stability can be estimated in around one minute has important bearings for neuropsychologists who must often perform testing sessions under strong time-constraints (Fig. 5).

**Figure 5:**
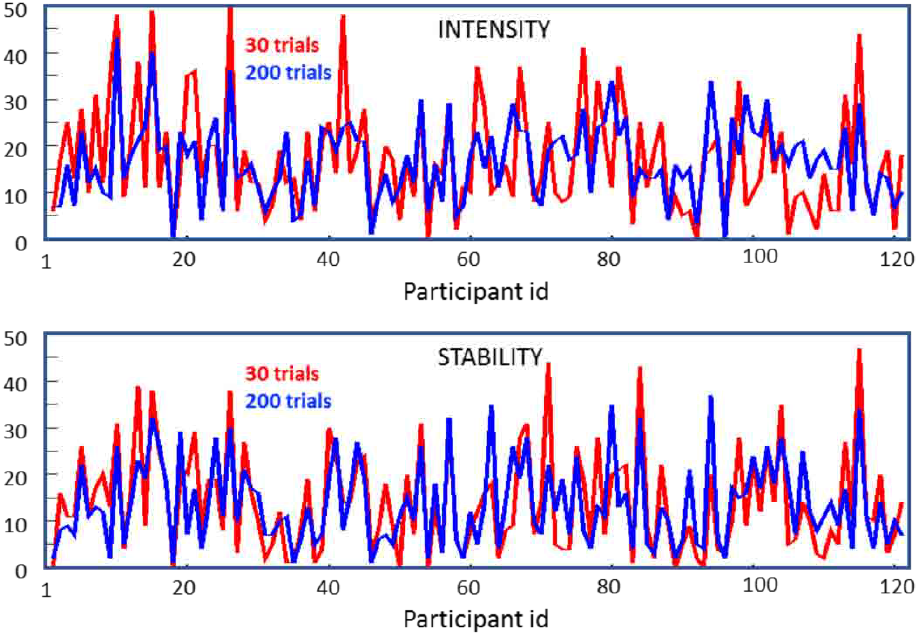
Comparison of INTENSITY and STABILITY computed over 30 vs. 200 trials. The graphs show values computed over 30 trials (red) versus 200 trials (blue) for INTENSITY and STABILITY for each participant.

**Table 3:** Correlation between behavioral indices measured over 200 trials, and the first 30, 40, 50, 60 trials of BLAST respectively (Pearson’s correlation coefficients).

|  | 30 trials | 40 trials | 50 trials | 60 trials | 200 trials |
| --- | --- | --- | --- | --- | --- |
| INTENSITY | 0.75 | 0.81 | 0.86 | 0.88 | 1 |
| STABILITY | 0.58 | 0.68 | 0.78 | 0.81 | 1 |
| FOCUS | 0.53 | 0.62 | 0.67 | 0.71 |  |
| MEAN_DURATION | 0.75 | 0.81 | 0.84 | 0.86 |  |
| MEDIAN_RT | 0.87 | 0.87 | 0.90 | 0.92 |  |

#### 4.2. Electrophysiological Study

This section describes the large-scale cortical dynamics during individual trials of BLAST and its variants, from the onset of the target letter to the motor response, with a millisecond and millimetric precision. Our strategy was to identify all major brain regions supporting BLAST from iEEG HFA_[50-150Hz]_ data and examine the specific dynamic of activation of each of those Regions of Interest (ROI) from iEEG of individual patients. The main objective was to associate precise cortical regions with each of the three main cognitive components of BLAST: i) encoding and maintenance of the Target letter, ii) visual search of the Array, iii) motor response. For that purpose, we compared HFA induced by BLAST and variants of BLAST lacking one or several of those components (C-BLAST and S-BLAST). We report only ROIs where similar activation patterns were observed in at least 2 different patients, following our standard procedure for iEEG analysis (Lachaux *et al*., 2012). Note that cortical regions with unspecific responses (the same response in all conditions) are not discussed here: which excludes for instance regions of the visual cortex that responded equally to all visual stimuli. Also, our analysis does not include subcortical responses, which might play an important role in BLAST but were not recorded. Finally, all sites with pathological activity and/or within the seizure onset zone were excluded from the analysis.

We found only seven ROIs with increased HFA during BLAST relative to baseline (Wilcoxon, p<0.01) (Fig. 6): the left Inferior Temporal Gyrus (ITG), the left ventral PreMotor Cortex (vPMC), the right and left Inferior Frontal Gyrus (IFG), the right dorsolateral PreFrontal Cortex (dlPFC), both left and right Frontal Eyes Field (FEF), a region anterior to the Supplementary Motor Area and immediately adjacent to the dorsal Cingulate Gyrus (preSMA) and the IntraParietal Sulcus bordering the Superior Parietal Lobule (IPS/SPL). In addition, five ROIs showed a reverse pattern of HFA *decrease* during BLAST: the middle Temporal Gyrus in the Lateral Temporal Cortex (MTG), the OrbitoFrontal Cortex (OFC), the ventromedial PreFrontal Cortex (vmPFC), the Temporal Parietal Junction (TPJ) and the Posterior Cingulate Cortex (PCC), all bilaterally.

**Figure 6:**
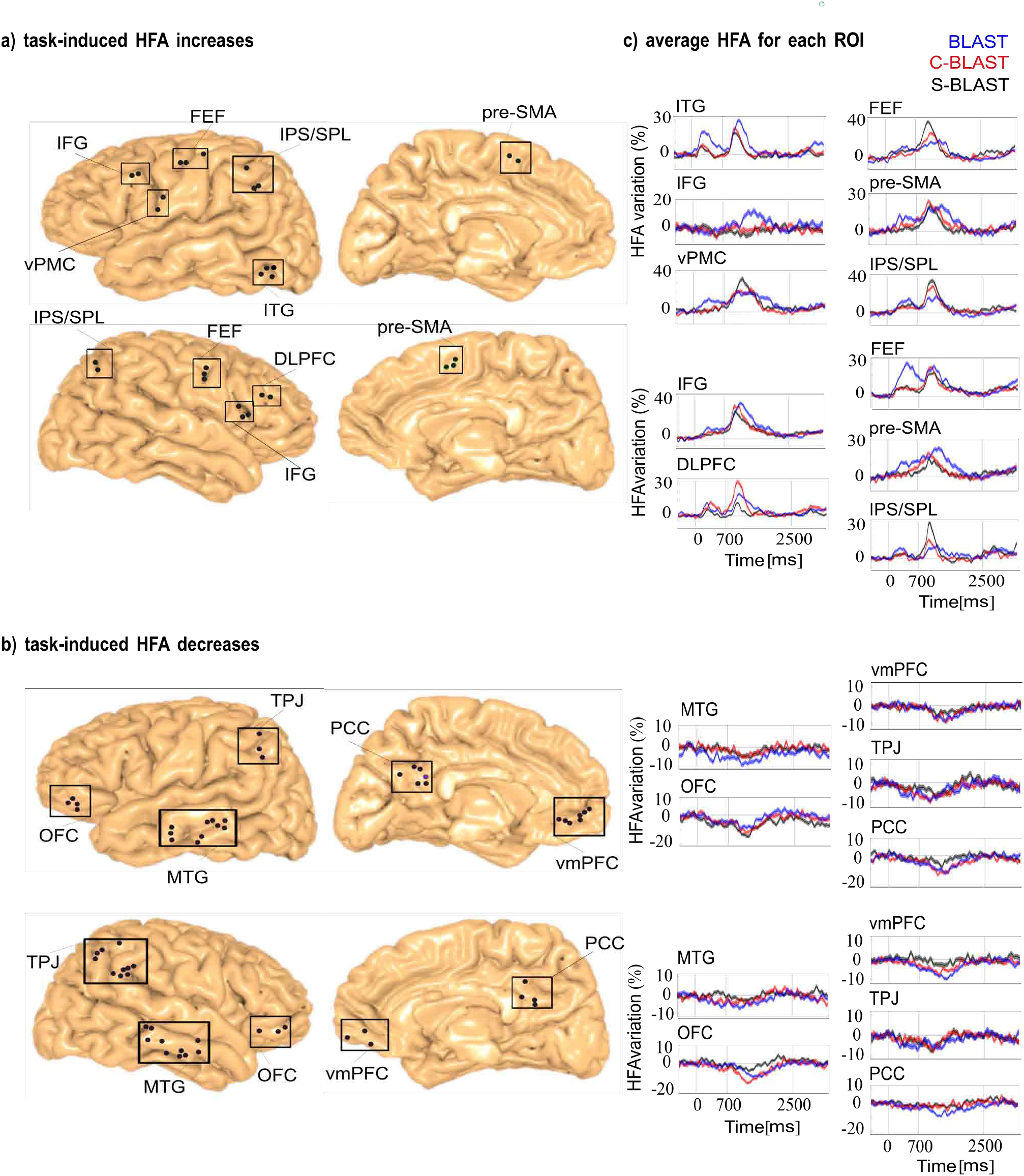
Cortical regions activated and deactivated during BLAST. a) anatomical location of task-related HFA increases during BLAST across all patients (each dot correspond to one particular SEEG recording site in a particular patient, non responsive sites are not shown). Mni-coordinates of individual iEEG sites are projected upon the MNI single-subject brain template; b) anatomical location of task-related HFA decreases. c) average dynamics of responsive sites in each ROI, expressed in % of the mean HFA value across the entire experiment. Vertical lines indicate cue-onset, cue-offset and array onset (from left to right). Experimental conditions are color-coded : blue for BLAST, red for C-BLAST, black for S-BLAST. Activated ROIs are the left inferior temporal cortex (ITG), the left ventral premotor cortex (vPMC), the inferior frontal gyrus (IFG), the right prefrontal cortex (PFC), the frontal eye fields (FEF), the dorsomedial prefrontal cortex (dmPFC) and the intraparietal sulcus/superior parietal lobule (IPS/SPL). Deactivated ROIs are the lateral temporal cortex (LTC), the orbitofrontal cortex (OFC), the ventromedial prefrontal cortex (vmPFC), the inferior parietal cortex (IPL) and the posterior cingulate cortex (PCC).

The portion of the left ITG active during BLAST matched the location and functional specificity of the word-form area, WFA, a region of the left inferior temporal lobe supporting visual processing of letters and letter-strings (Fig. 7). The dynamics of activation during BLAST revealed two phases: during the presentation of the Target and during the display of the Array, as expected in response to letters. However, the statistical comparison of BLAST and its variants C-BLAST and S-BLAST revealed that the response to the letter was strong and sustained only when the Target must be encoded into working memory (BLAST) (Fig. 7). It was also stronger when the Array was searched attentively (i.e. in BLAST). Therefore, the WFA participated actively to both the encoding phase and the search phase of BLAST.

**Figure 7:**
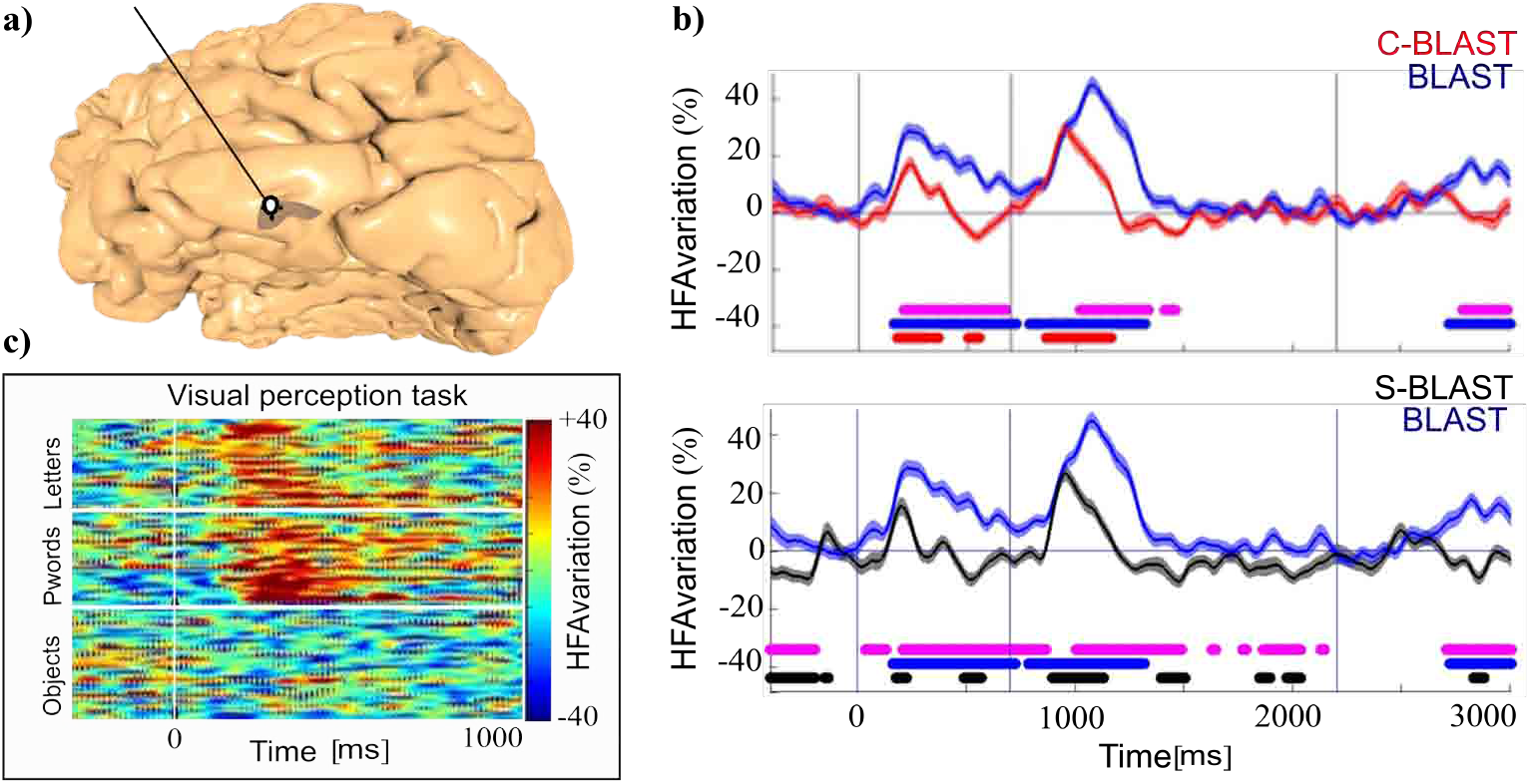
Example of individual response in the left ITG during BLAST. a) Exact location of the recording site reconstructed onto the patient’s individual 3D MRI. b) HFA increase during BLAST (blue), C-BLAST (red) and S-BLAST (black) expressed in % of the average HFA across the entire experiment for that site. Horizontal lines indicate time windows with a significant deviation relative to the pre-stimulus baseline level in each condition separately (same color code as above). Pink horizontal lines indicate time windows with a significant difference between the two conditions in the graph. c) HFA increase induced by flashed pictures of different categories (object, pseudo-words and consonant strings) during a visual oddball task performed by the patient in a separate session. Picture onset is at 0 ms, and each horizontal line corresponds to one single picture. The matrix illustrates the specificity of the visual response to letter strings, for that site (MNI coordinates, −66 −49 −17).

In the inferior frontal cortex, the left IFG was only active during the Array presentation and selectively in the BLAST task, with a stronger activity when the participant took longer to respond (Fig. 8). In contrast, the left vPMC and the right IFG reacted to both the Target and the Array, with a stronger response in the BLAST condition (Fig. 8, and 9). Note that the letter stimulus has the same physical characteristics in BLAST, C-BLAST and S-BLAST, but carries task-relevant information only in BLAST, in which it must be encoded and maintained in short-term memory to serve as a template for the visual search. Single trial representation (matrix representation in Fig. 9) shows that activity was sustained continuously until task completion in the IFG, but extended beyond the reaction time in the vPMC, suggesting a participation to motor programming and execution for that region rather than to the more cognitive aspects of BLAST.

**Figure 8:**
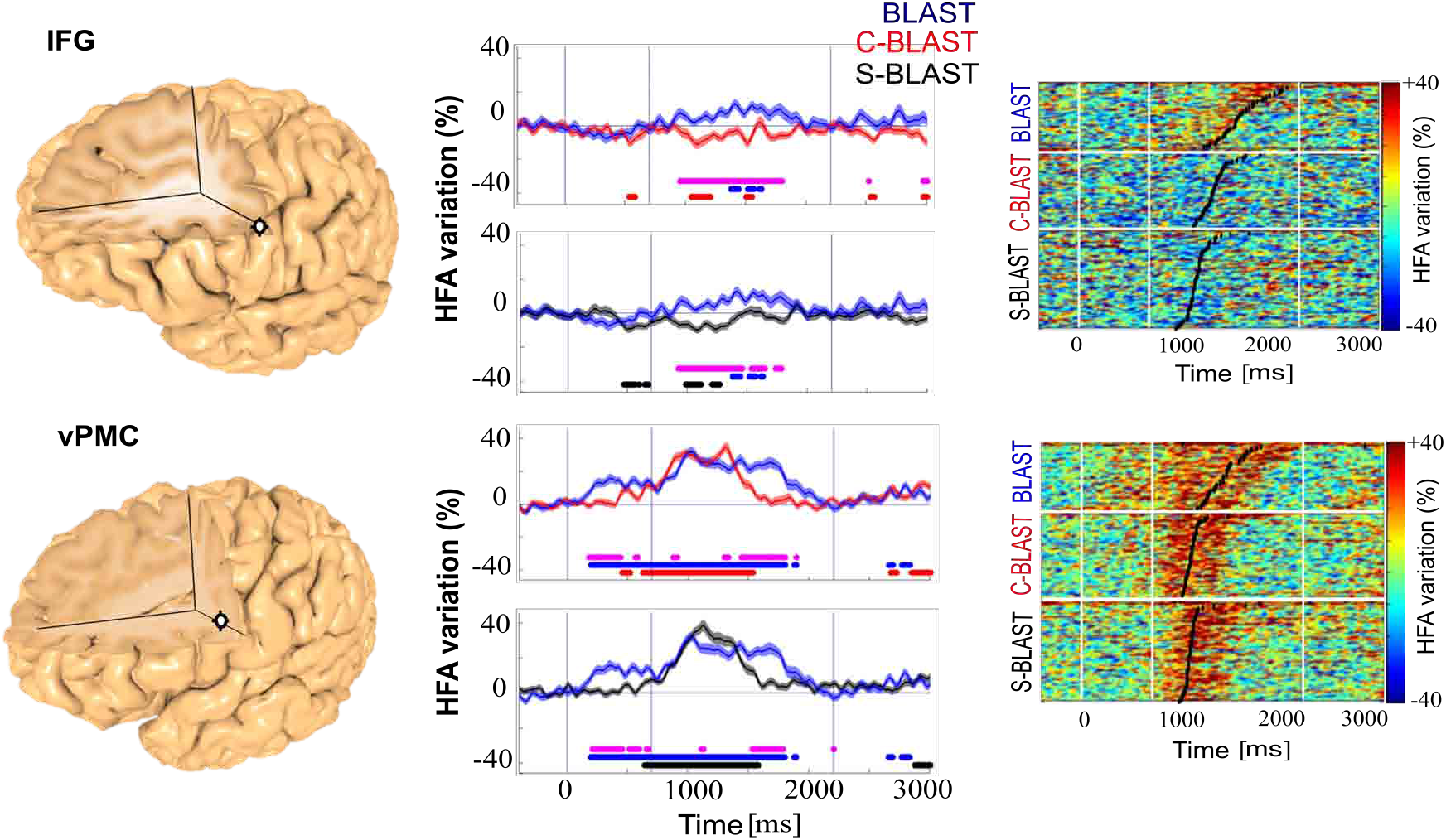
Example of individual response in the left IFG and left vPMC during BLAST. Same legend as fig. 7. Response to the Array in the left IFG (top) was especially strong in the BLAST condition. MNI coordinates for this site were [−57 19 24]. Response in the left vPMC (bottom) was especially strong to the Target in the BLAST condition. MNI coordinates for this site were [−49 −1 21]. (See Supplementary Fig.3 for more examples).

**Figure 9:**
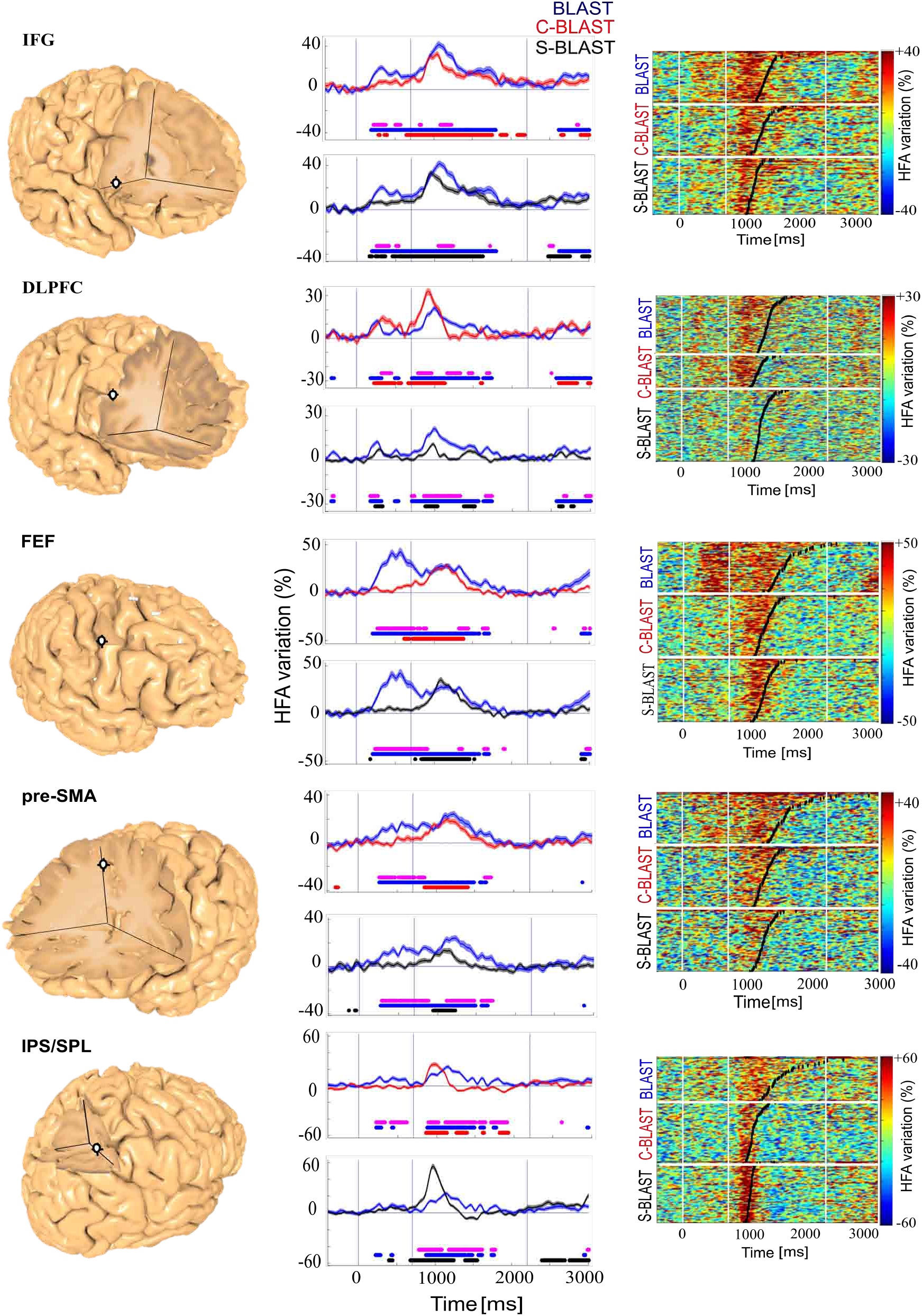
Example of individual response in the right fronto-parietal cortex. Same legend as fig. 7. Responses were especially strong to the Target in the BLAST condition for all sites. MNI coordinates: in right IFG [50 21 4]), right DLPFC [50 33 14], right FEF [54 1 34], right pre-SMA [11 10 48] and right IPS/SPL [29 −58 44]). A similar pattern was also observed in the left FEF [−37 −11 51], left pre-SMA [−10 14 52] and left IPS/SPL [−41 −45 38]. (See Supplementary Fig.3).

All other ROIs stopped their activity upon task completion. The activation of the superior parietal lobe (IPS/SPL) occurred only during the Array with a stronger response in the two BLAST variants with a red singleton in that Array (Fig. 9), compatible with a role of that region in spatial attention shifts and attentional capture by salient stimuli. Activation during Target encoding in BLAST indicated that it might participate in a large-scale top-down process to sustain activation in letter-specific areas of the inferior Temporal Lobe. Such top-down process have been modelled within a larger Dorsal Attention Network (Corbetta and Shulman, 2002), which also includes the Frontal Eye Field, or FEF (Fig. 9): accordingly, we found a region highly reactive to the Array in the lateral motor cortex around the putative FEF location. Interestingly, single subject analysis revealed a specific response in that region to Targets which must be encoded into working memory (BLAST), and must therefore be attended.

Similar patterns of activation were observed in the right dorso-lateral PFC and the preSMA (Fig. 9), immediately adjacent to the dorsal anterior cingulate gyrus. Both regions are hallmarks of the frontal executive system which mediates cognitive control and goal-directed behavior in general, and the ability to stay-on-task in particular. Unsurprisingly, both ROIs were active during the encoding and during the search phase, with a steady activity increase in the latter until task completion.

In the five ROIs deactivated by BLAST, HFA suppressions observed in individual patients revealed a stronger deactivation in the most difficult task (BLAST) in most instances (Fig. 10). They had a slow dynamics with a negative peak while participants processed the Array. Single trial matrixes suggest that the five ROIs might have different temporal characteristics; but HFA decreases, although significant, were not sufficiently strong to elaborate further on their dynamics, in contrast with the massive HFA increases seen in other ROIs. All ROIs deactivated during BLAST are part of the Default-Mode Network (Raichle *et al*., 2001), in line with recurrent observations that activity is suppressed in the DMN when processing external stimuli attentively.

**Figure 10:**
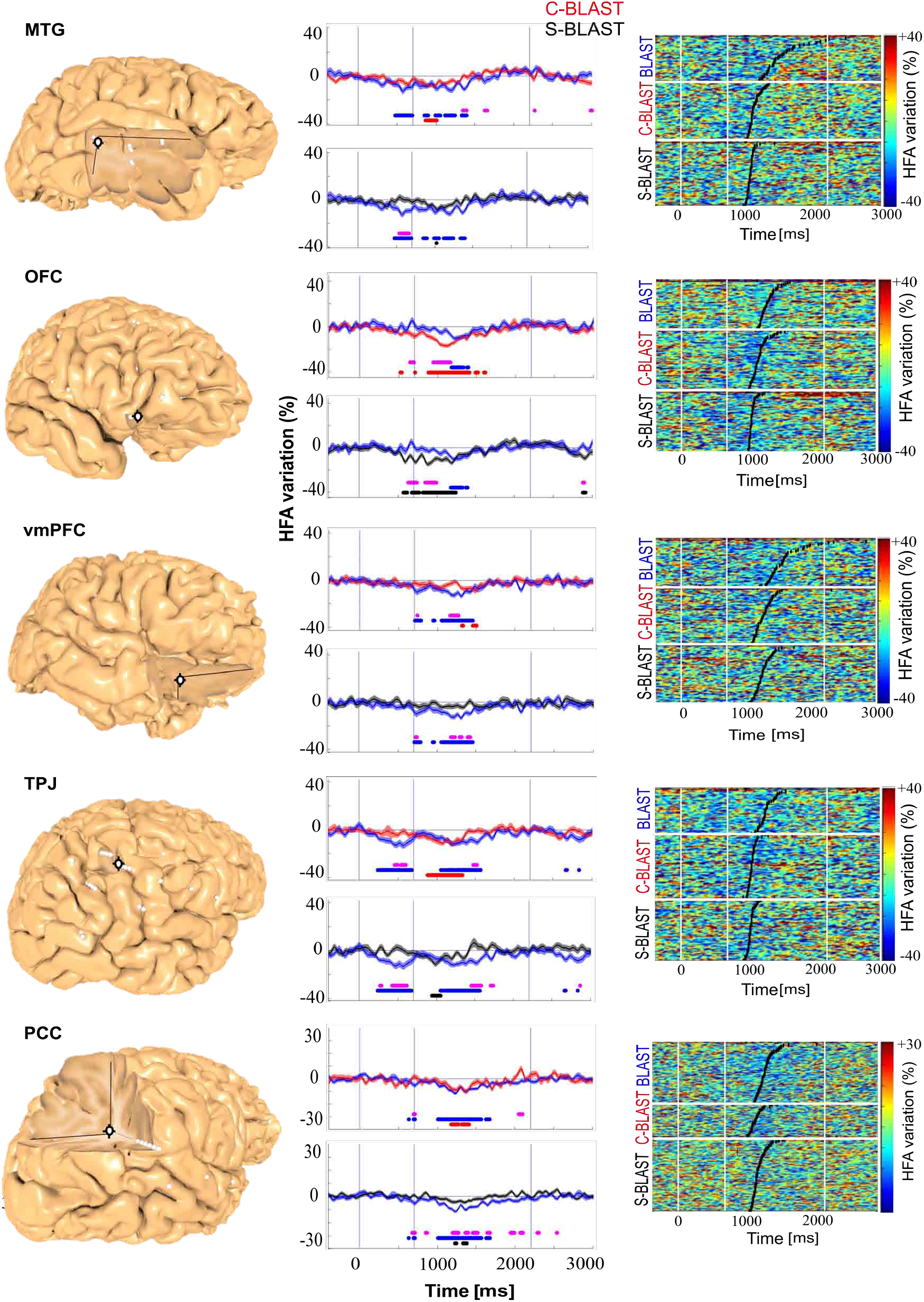
Example of neural deactivations in the right hemisphere. Same legend as in fig.7. The deactivation was especially strong to the Target in the BLAST conditions for sites registered in the right MTG (MNI coordinate [57 −31 −3]) and right TPJ (MNI coordinate [51 −44 50]). The deactivation was especially strong to the Array in the BLAST condition in right OFG (MNI coordinate [47 45 −11]), right vmPFC (MNI coordinate [7 27 −26]) and right PCC (MNI coordinate [11 −46 29]). A similar pattern was also observed in the left sites (see Supplementary Fig.4 for more examples).

## 5. Discussion

Our objective was to propose a test that measures « the ability to stay on-task », with a well-identified cortical network and age-stratified normative data. We focused on the ability to apply consistently and selectively the set of cognitive processes necessary and sufficient to perform the task at hand, with no interference from irrelevant processes. BLAST (Bron/Lyon Attention Stability Test) is to our knowledge the first test meeting all such requirements, with a performance that is related selectively to inattention, not hyperactivity. In addition, intracranial EEG data provide clear evidence that BLAST involves primarily the frontal executive and dorsal attention network, including all major components of the executive attention system in the prefrontal cortex, together with a reduction of activity in the Default-Mode Network (often termed “task-negative network”, Fox *et al*., 2005).

### 5.1. Task design

We argue here that there are very few possible alternatives to BLAST, in terms of task-design, to measure attentional stability with a close-to-second temporal resolution. BLAST provides a reaction time every two seconds on average, which defines the temporal precision at which fluctuations of attention can be detected. It matches the time-scale of short MLA which can make us « miss » an important word in an explanation, for instance. It is hard to imagine a task with a better temporal precision, considering the minimal duration of cognitive stimulus-to-response cycles (Madl *et al*., 2011) and the constraint to avoid muscular fatigue. In theory, a finer time resolution could be achieved if participants had to react continuously to a changing stimulus (with a continuous behavioral response, as when tracking a moving dot with a joystick); however, to the best of our knowledge, no such task keeps difficulty constant over time (for instance, in tracking tasks, parts of the trajectory with high, changing curvatures are harder to track than smoother parts), and therefore, performance over time does not depend solely upon the participant’s attention. Tasks in which behavioral measures are discrete in time, like BLAST, involve repeated presentations of a sensory stimulus followed by a participant’s overt response, at a pace which determines the temporal resolution of the performance measurement. Such discrete tasks must themselves obey several constraints to ensure that performance depends solely (or as much as possible) upon the attention allocated by the participant to the task: i) the inter-stimulus interval should be constant (to keep task difficulty constant) and on the order of one or two seconds to avoid muscular fatigue and allow reasonable time to process the stimulus; ii) the rule defining stimulus-response association should vary from trial to trial to minimize the possibility of automatization (which would reduce the behavioral impact of lapses of attention) : this implies that participants should be informed of that association-rule before each trial by a cue, shown before stimulus presentation so that cue- and stimulus-processing do not compete for attention resources; and yet iii) the overall task-set (the principle of stimulus-response association) should remain the same throughout the task, as it is known that task-set changes within tasks induce switch costs which affect behavior negatively independently of attention. This set of constraints sums up to allow only designs in which in each trial, a cue instructs participants - with a metronomic timing - to process the upcoming stimulus in a unique, but yet stereotyped way, to choose one of several motor responses. In BLAST, the cue is a target letter which instructs participants to decide whether that letter is among the four letters shown 700 ms after that stimulus. The stimulus-response association rule is not rigid (the same stimulus is not always associated with the same motor response), and yet, the task set remains the same throughout the task. Alternatives to BLAST are of course possible to track attention on a second-to-second basis, but we claim that all of them would follow the same organizing principle.

There are of course computerized tests to evaluate the ability to sustain attention over time and help diagnose ADHD, such as the Conners’ Continuous Performance Test (Homack and Riccio, 2006). But none of them was designed to capture brief lapses of attention as only a minority of the trials really require attentive processing. Interestingly, a task-design very similar to BLAST has been used to study the neural mechanisms of visual attention in non-human primates: the task used an array of four tilted bars instead of letters (Buschman and Miller, 2007). But the trials did not repeat as in BLAST and the authors did not study the variability of reaction times.

### 5.2. Neural correlate of BLAST

Our electrophysiological study suggests the following scenario during successful trials of BLAST: after an initial and transient response of early visual areas, activation triggered by the Target letter would reach a region of the basal temporal lobe specialized for letter-forms: the Word Form Area or WFA. The WFA would then hold the search template until and throughout the search process (the display of the Array). If the participant strategy implies subvocal rehearsal, the Inferior Frontal Gyrus would support an additional, phonological, maintenance process using the phonological loop (in agreement with the classic model of verbal working memory (Baddeley, 2000). The Dorsal Attention Network, or DAN (Corbetta and Shulman, 2002), including the dorsolateral Prefrontal Cortex, the Frontal Eye Field and the Intraparietal sulcus would a) facilitate visual memory maintenance in the WFA (as the DAN has been shown to sustain and bias activity in visual areas via top-down influences, Noudoost *et al*., 2010; Seidl *et al*., 2012), and b) guide visual attention within the Array in line with its primary functional role (Corbetta and Shulman, 2002). The DLPFC might orchestrate the encode-then-search process, given its known ability to maintain task-instructions (the “task sets”, Sakai, 2008)) : it would hold the « program » to encode the search template (the Target) in the WFA/IFG, then compare sequentially or in parallel each letter of the Array with that template and support the decision process to press with the right or left index finger. Finally, a region at the interface between the pre-SMA and the dorsal Anterior Cingulate Gyrus would ensure that attention is « on-task », in line with its known implication in cognitive control. The final motor response would obviously be produced by the premotor and motor cortex. In that scenario, a sudden increase in reaction times might be due to a less efficient encoding and maintenance of the search template in the WFA (insufficient top-down influences from the DAN), or a discontinuous search process (again, inefficient or insufficient activation of the DAN or the pre-SMA).

This global and highly organized activation pattern does not occur in isolation, but together with a somewhat smoother deactivation process of an entire network called DMN (the Default-Mode Network or task negative network: the Temporal Parietal Junction, the posterior Cingulate Cortex, the medial Prefrontal Cortex, the ventral lateral Prefrontal Cortex and the lateral Temporal Cortex) (Raichle *et al*., 2001). Activity in the DMN is classically associated with spontaneous, task-unrelated cognition, which might interfere with active visual processing of external stimuli, and its deactivation during BLAST was predicted by the literature (Weissman *et al*., 2006; Li *et al*., 2007; Esposito *et al*., 2009; Anticevic A1, Repovs G, Shulman GL, 2010; Mayer *et al*., 2010; Anticevic, 2013). However, the precision of iEEG reveals a surprisingly dynamic behavior with a brief reactivation in the ultra-short interval (800 ms) between two consecutive trials: a brief attempt to reactivate DMN functions whenever possible (Lachaux *et al*., 2008; Ossandon *et al*., 2012).

To summarize, BLAST exemplifies the interplay between the DAN and the DMN which is typical of sustained and demanding visual attention tasks (Langner and Eickhoff, 2013; Christoff *et al*., 2016). It is therefore ideal to study the ability to stay on-task, and follow the dynamic competition between the DAN and the DMN on a trial-by-trial basis, with second-by-second precision.

A first possible objection to the present study is that the cortical network supporting BLAST was identified in patients suffering from epilepsy. Although it is true that their brain can only be considered as a proxy of a healthy brain, that criticism has been addressed repeatedly over the years and detailed responses including precise guidelines have been published (see for instance, Lachaux et al., 2012). In short: researchers analyzing iEEG carefully check that recordings are made outside epileptogenic networks and are free of epileptiform activity; and that similar observations can be made in the same cortical structure in patients with different types of epilepsy and medications. Thanks to such sanity checks, iEEG has become in twenty years a premium methodology to study the neural dynamics of human cognition (Lachaux et al., 2012; Ritaccio et al., 2014), which results are now widely incorporated in our global understanding of human brain functions. Regarding the task design, one might criticize that letters used in BLAST can be verbalized. But that would be true of any familiar symbol (such as geometric shapes like squares or circles or ideograms). With random, unknown shapes, we could not exclude that participants familiarize with the symbols over time, making the task eventually easier. We chose letters, because they were arguably the most familiar symbols to almost any French individual older than 6. A second remark is that BLAST difficulty is not strictly constant across trials, because some letters might be easier to spot than others, because the location of the target within the array (when present) is likely to have an impact, and because it might take longer to figure out that the target is not in the array. We acknowledge such effects fully, but argue - and checked visually - that the variations in reaction times induced by such factors were minimal compared to the effect of momentary lapses of attention. A third possible criticism changes of strategy might cause fluctuations of reaction times independently of attention (for instance, in a change in the speed/accuracy trade-off), but that particular criticism applies to any design using reaction times to assess attention. Our assumption is that changes of strategy should not cause instability of reaction times on a trial-to-trial basis, but rather a slower change with stable periods lasting several trials; therefore, it is likely that short-scale fluctuations in reaction times that we take as a measure of inattention are indeed due to inattention. Finally, we are fully aware that behavioral performance can only provide an indirect estimate of the level of attention allocated to the task; however, one might argue that in real-life situations, it is also performance, not attention in itself, which ultimately matters.

### 5.3. Possible limitations of BLAST

A first possible objection to the present study is that the cortical network supporting BLAST was identified in patients suffering from epilepsy. Although it is true that their brain can only be considered as a proxy of a healthy brain, that criticism has been addressed repeatedly over the years and detailed responses including precise guidelines have been published (see for instance, Lachaux *et al*., 2012). In short: researchers analyzing iEEG carefully check that recordings are made outside epileptogenic networks and are free of epileptiform activity; and that similar observations can be made in the same cortical structure in patients with different types of epilepsy and medications. Thanks to such sanity checks, iEEG has become in twenty years a premium methodology to study the neural dynamics of human cognition (Lachaux *et al*., 2012; Ritaccio *et al*., 2014), which results are now widely incorporated in our global understanding of human brain functions.

Regarding the task design, one might criticize that letters used in BLAST can be verbalized. But that would be true of any familiar symbol (such as geometric shapes like squares or circles or ideograms). With random, unknown shapes, we could not exclude that participants familiarize with the symbols over time, making the task eventually easier. We chose letters, because they were arguably the most familiar symbols to almost any French individual older than 6. A second remark is that BLAST difficulty is not strictly constant across trials, because some letters might be easier to spot than others, because the location of the target within the array (when present) is likely to have an impact, and because it might take longer to figure out that the target is not in the array. We acknowledge such effects fully, but argue - and checked visually - that the variations in reaction times induced by such factors were minimal compared to the effect of momentary lapses of attention. A third possible criticism changes of strategy might cause fluctuations of reaction times independently of attention (for instance, in a change in the speed/accuracy trade-off), but that particular criticism applies to any design using reaction times to assess attention. Our assumption is that changes of strategy should not cause instability of reaction times on a trial-to-trial basis, but rather a slower change with stable periods lasting several trials; therefore, it is likely that short-scale fluctuations in reaction times that we take as a measure of inattention are indeed due to inattention. Finally, we are fully aware that behavioral performance can only provide an indirect estimate of the level of attention allocated to the task; however, one might argue that in real-life situations, it is also performance, not attention in itself, which ultimately matters.

### 5.4. Conclusion

In summary, BLAST provides one of the few possible task designs to measure fluctuations of executive attention behaviorally on a second-to-second basis, and because performance is measured so often, it can provide a first indication of an individual’s ability to stay on task in less than a minute, with numerous possible applications (testing the effects of a pharmacological treatment, evaluating the cognitive impact of short pathophysiological events such as epileptic spikes, measuring the benefits of cognitive rehabilitation or attention training programs,…). In addition, the comparison of performance between BLAST and its simplified versions can potentially reveal a deficit in a specific cognitive process, such as encoding/maintenance or visual search, which can then be related with specific brain regions thanks to our detailed knowledge of the BLAST cortical network. For basic research purposes, BLAST is also currently used to understand the neural basis of endogenous attention fluctuations and MLA, and the neural correlates of transient performance decrement observed after external distractions or during multi-tasking.

## 1. Supplementary figures

**Sup Figure 1:**
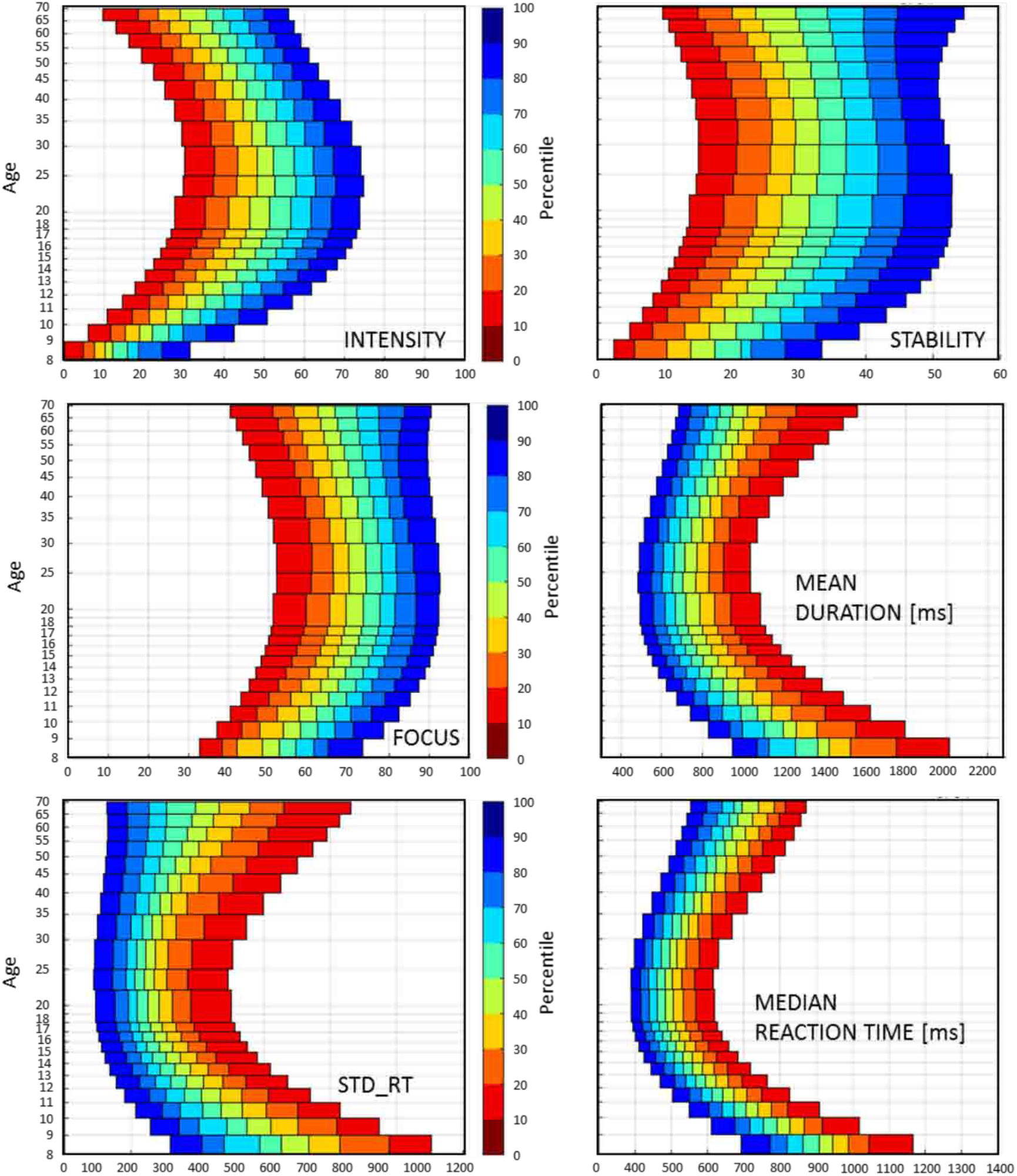
Models of Age-stratified normative data for the main behavioral indices quantifying the ability to stay on task in BLAST Third-order Polynomial best-fits of the distributions.

**Sup Figure 2 :**
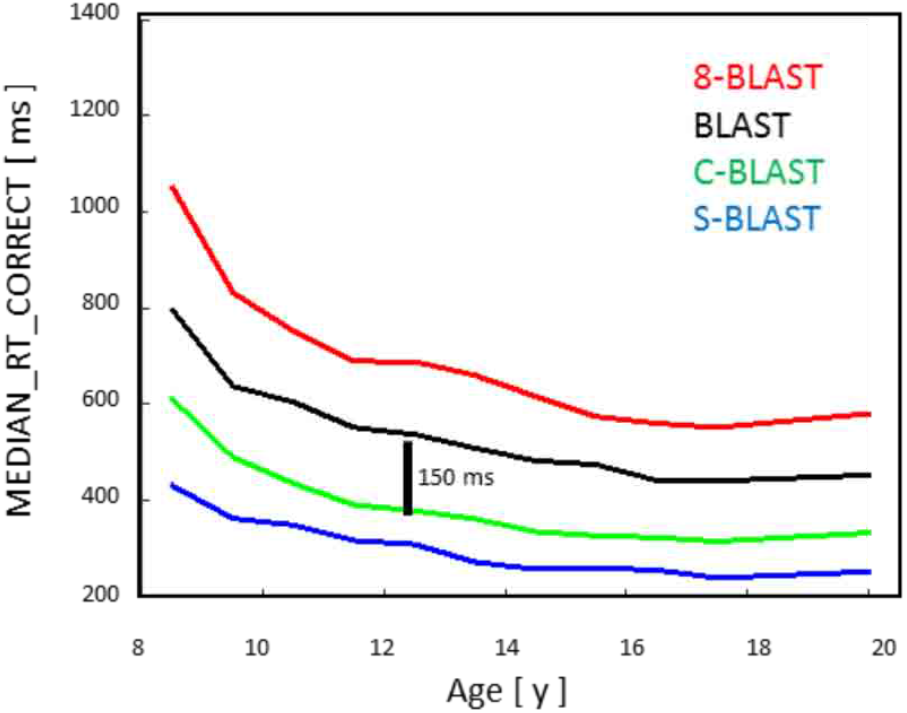
Median reaction time for each of the adaptive versions of BLAST.

**Sup Figure 3 :**
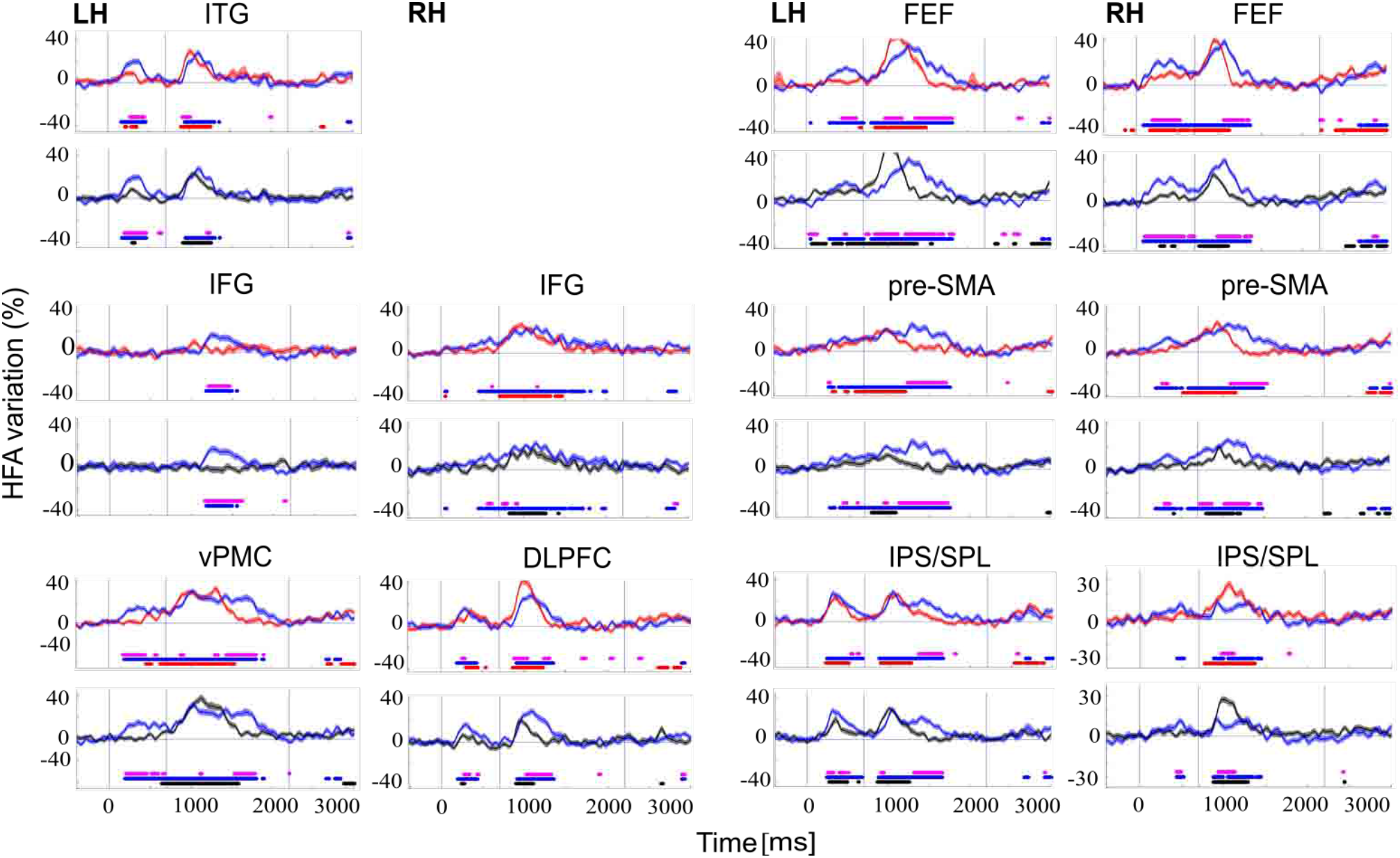
Additional examples of individual ROI activations during BLAST. HFA [50-150Î] increase during BLAST (blue), C-BLAST (red) and S-BLAST (black) expressed in % of the average HFA across the entire experiment for that site. Horizontal lines indicate time windows with a significant deviation from the pre-stimulus baseline level in each condition separately (same color code as above). Pink horizontal lines indicate time windows with a significant difference between the two conditions in the graph. Target onset is at 0ms. MNI coordinates for each site: Left Hemisphere (LH) : ITG [−53 −47 −15], IFG [−48 18 28], vPMC [−53 0 14], FEF [−37 −11 51], preSMA [-10 14 52] and IPS/SPL [−29 −34 42]); Right Hemisphere (RH): IFG [44 13 10], DLPFC [36 29 16], FEF [47 0.5 31], preSMA [12 15 45] and IPS/SPL [29 −59 44]).

**Sup Figure 4 :**
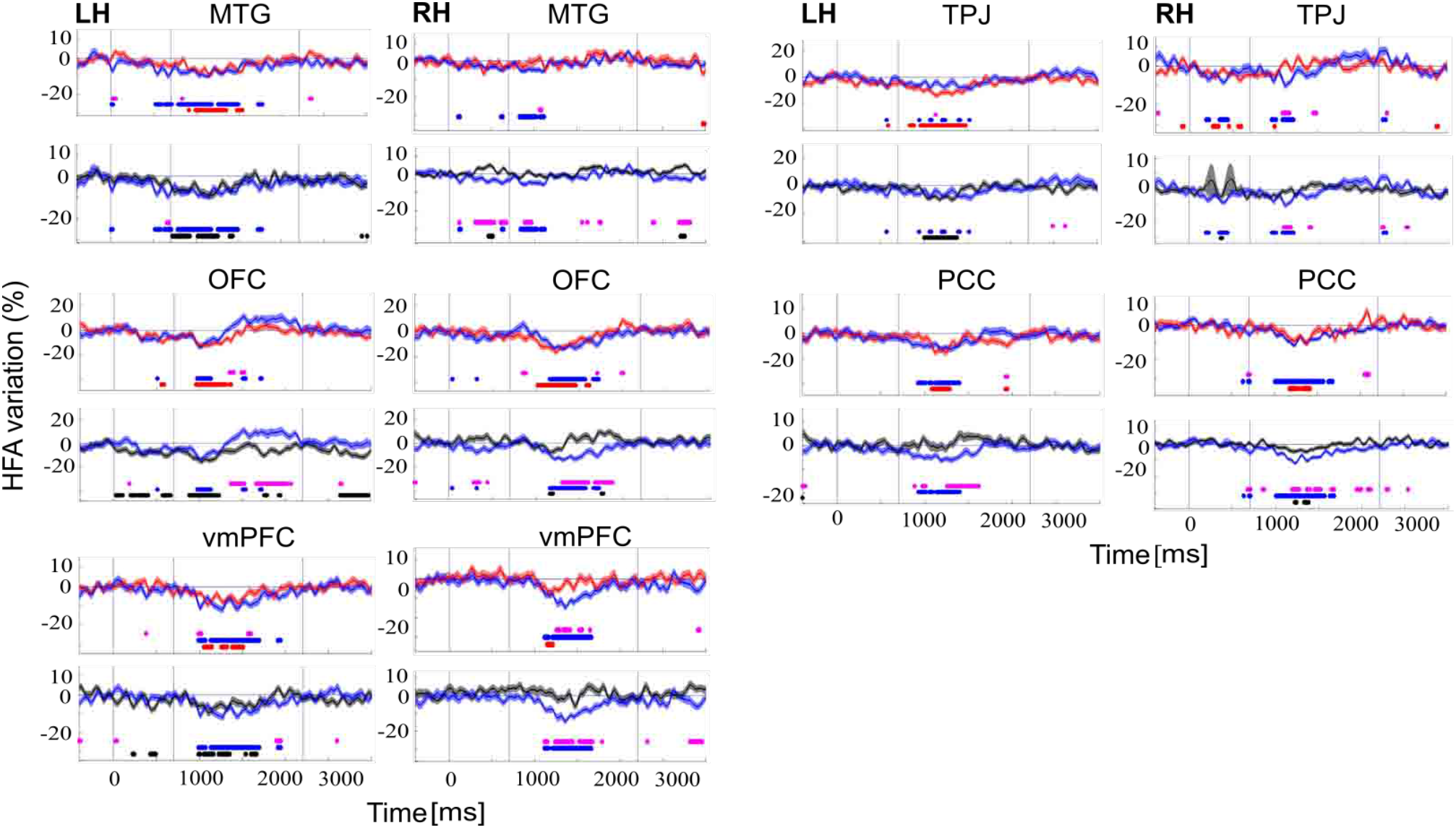
Additional examples of individual deactivations in BLAST ROI Same legend as supplementary fig. 3. MNI coordinates for each site : Left Hemisphere: MTG [−55−14 −16], OFC [−36 42−12], vmPFC [−8 53 −6], TPJ [−44 −46 56] and PCC [−8 −58 30]); Right Hemisphere: LTC [63 −31 −13], OFC [36 40 −14], vmPFC [9 40-14], TPJ [59 −35 32] and PCC [10 −35 29]).

